# ATM Inhibition Synergizes with Fenofibrate in High Grade Serous Ovarian Cancer Cells

**DOI:** 10.1101/2020.05.29.123919

**Authors:** Chi-Wei Chen, Raquel Buj, Erika S. Dahl, Kelly E. Leon, Katherine M. Aird

## Abstract

**Background:** Epithelial ovarian cancer (EOC) is the deadliest gynecological malignancy in the United States with high grade serous ovarian cancer (HGSOC) as the most commonly diagnosed subtype. While therapies targeting deficiencies in the homologous recombination (HR) pathway are emerging as the standard treatment for HGSOC patients, this strategy is limited to the 50% of patients with a deficiency in this pathway. Therefore, patients with HR-proficient tumors are likely to be resistant to these therapies and require alternative strategies.

**Methods:** Data from HGSOC patients in The Cancer Genome Atlas (TCGA) were analyzed for ATM status, ATM and PPARα expression, and used to perform Gene Set Enrichment Analysis (GSEA). Screening data from the Dependency Map were analyzed to identify FDA-approved drugs that preferentially inhibit ATM-low cancer cells. *In vitro* studies were performed to determine whether ATM inhibitors synergize with the PPARα agonist fenofibrate in HGSOC cell lines.

**Results:** The HR gene Ataxia Telangiectasia Mutated (ATM) is wildtype in the majority of HGSOC patients and its kinase activity is upregulated compared to normal fallopian tube tissue. As high ATM has been associated with poor overall and progression-free survival, targeting ATM may be beneficial for a subset of HGSOC patients. Clinical trials of ATM inhibitors are commencing; however, ATM inhibitors are not effective as single agents. We aimed to explore novel therapeutic vulnerabilities of ATM deficient cells to develop a combinatorial therapy. Using data from TCGA, we found that multiple pathways related to metabolism are inversely correlated with ATM expression, suggesting that combining ATM inhibition and metabolic inhibition would be effective. Indeed, analysis of FDA-approved drugs from the Dependency Map demonstrated that ATM low cell lines are more sensitive to fenofibrate, a PPARα agonist that has been previously shown to affect multiple cellular metabolic pathways. Consistently, PPARα signaling is associated with ATM expression. We validated that combined inhibition of ATM and treatment with fenofibrate is synergistic in multiple HGSOC cell lines by inducing senescence.

**Conclusions:** Our results suggest that metabolic changes induced by ATM inhibitors are a potential target for the treatment for HGSOC.

## Background

Epithelial ovarian cancer (EOC) remains the most lethal gynecological malignancy (1). EOCs are divided into multiple subtypes with high grade serous ovarian cancer (HGSOC) as the most common. Most HGSOC patients are diagnosed at advanced stages (III-IV), and the 5-year survival rate for these patients is <30%. Current standard-of-care for HGSOC is debulking surgery followed by platinum-based chemotherapy (2). While the majority of patients initially respond to therapy, relapse with chemoresistant disease occurs in a significant number of patients (3). Recurrent disease is treated with poly(ADP)ribose polymerase (PARP) inhibitors if the patient is BRCA mutant or previously responded to platinum-based therapy (4). Response to platinum and PARP inhibitors occurs due to deficiencies in the DNA damage repair pathway homologous recombination (HR) (5). Homologous recombination deficiency (HRD) occurs in ~50% of patients and includes loss-of-function mutations in multiple proteins related to HR-mediated DNA repair (4). Unfortunately, the 50% of HGSOC patients with HR-proficient disease typically do not respond well to current therapies and have worse overall survival (6). Therefore, identification of therapies to treat this subset of HGSOC patients is urgently needed.

Ataxia Telangiectasia Mutated (ATM) is a serine/threonine kinase that is critical for HR-mediated repair of DNA double strand breaks (DSBs) (7,8). Upon sensing of DSBs by the MRN complex, ATM is recruited to the damage site, where it is first activated by autophosphorylation of S1981 (9). Upon activation, ATM phosphorylates a number of downstream substrates, including Chk2 and H2AX, that promote cell cycle arrest/delay and DNA repair (8). Germline mutations in ATM lead to the disorder Ataxia Telangiectasia (A-T), which has a number of pathological consequences, including predisposition to cancer and metabolic dysfunction (7,8). Given that A-T patients have a higher risk of cancer and Atm−/− mice develop malignancies (7,8,10), ATM has been thought to be a tumor suppressor. However, many tumors rely on elevated DNA repair pathways, and a recent publication demonstrates that ATM is required for tumorigenesis (11). Additionally, a previous study in HGSOC showed that patients with high nuclear ATM expression have worse survival (12). Together, these studies suggest that ATM may be an actionable target in the subset of tumors where it is wildtype and elevated. Indeed, ATM inhibitors have been in clinical development for the past two decades (13,14). Multiple pre-clinical studies have indicated that ATM inhibitor monotherapy is not likely to be effective (14–18). However, combined inhibition of ATM and DNA damaging agents such as PARP inhibitors and irradiation is synergistic (18), and recently a Phase I clinical trial using the ATM inhibitor AZD0156 in combination with a variety of DNA damaging agents has commenced (clinicaltrials.gov). As ATM inhibition also affects metabolic pathways (19–21), identifying targets beyond DNA damaging agents for combinatorial therapy with ATM inhibitors may therefore open up a new paradigm for treatment.

Peroxisome Proliferator Activated Receptors (PPARs: PPARα, PPARδ, PPARγ) are nuclear receptors and ligand-inducible transcription factors (22,23). Upon ligand induction, PPARs heterodimerize with retinoid X receptor (RXR) and drive transcription of a variety of downstream targets. Downstream targets of the PPARs differ greatly, likely due to dissimilarity in endogenous and exogenous ligands. Ligands for PPARα include endogenous ones like long chain unsaturated fatty acids and exogenous ones such as fibrates (23). Activation of PPARα leads to transcription of multiple metabolic genes, such as those related to fatty acid oxidation or inhibition of glycolysis (24). This has been therapeutically exploited in patients with dyslipidemia by using fenofibrate or clofibrate, exogenous ligands for PPARα (25). The role of PPARα in cancer is not fully defined, as it is tumor-promoting in rodents but not humans and tumor suppressive in a context- and cancer-type dependent manner (26). Activation of PPARα by treating cancer cells with fenofibrate or clofibrate alone or in combination with other drugs has been shown to decrease proliferation and survival through a variety of mechanisms (27). However, the combination of fenofibrate and ATM inhibitors has never been explored.

Here we show that ATM is wildtype and upregulated in HGSOC. Analysis of HGSOC TCGA datasets found that multiple metabolic pathways are associated with low ATM, suggesting that these pathways could be targeted in combination with ATM inhibitors. Using data from Dependency Map, we identified the PPARα agonist fenofibrate as an FDA-approved drug that inhibits ATM-low cell survival to a greater extent than ATM-high cancer cells. Indeed, PPARα correlates with ATM expression in HGSOC patient specimens. Consistent with high throughput data, combined inhibition of ATM and treatment with fenofibrate is synergistic in HGSOC cells by inducing senescence. These results provide a proof-of-principle study to used combined inhibition of ATM and treatment with a metabolic drug for HGSOC therapy.

## Methods

### TCGA database and GSEA analysis

Spearman’s correlation in HGSOC (TCGA, PanCancer Atlas) was calculated using cBioportal (28,29) for each gene in the RNA-Seq. Genes were ranked according to the Spearman correlation coefficient and the p-value of the correlation as follows: −log_10_(p value)*sign(log_2_ correlation coefficient) (30). Pre-ranked files were used to run pre-ranked GSEA (MSigDB collection KEGG and Reactome) (31) under predefined parameters. Following GSEA documentation: https://software.broadinstitute.org/cancer/software/gsea/wiki/index.php/FAQ#Why_does_GSEA_use_a_false_discovery_rate_.28FDR.29_of_0.25_rather_than_the_more_classic_0.05.3F. Terms with a q-value < 0.25 where considered significant.

### Dependency Map data analysis

Raw data from the PRISM drug repurposing screen (32) and reverse phase protein array (RPPA) (33,34) were downloaded from depmap.org. Cell lines were divided in half based on ATM protein expression. Only “launched” drugs were considered. Drugs were considered “hits” if log_2_foldchange <0 (ATM low vs. ATM high) and FDR <0.25. Other analyses were performed using the depmap.org online tool.

### Cell lines and culture conditions

Ovcar3 and Ovcar10 HGSOC cells were cultured in RPMI-1640 (Corning, Cat# 10-040-CV) supplemented with 5% FBS. All cell lines were cultured in MycoZap and were routinely tested for mycoplasma using a highly sensitive PCR-based method (35). Tumor cell lines were authenticated using STR Profiling using Genetica DNA Laboratories.

### Colony Formation

Cells (5×10^6^/well in 12-well plates) were seeded, allowed to adhere overnight, and washed twice with 1x PBS the next day. Cells were cultured with serum-free RPMI-1640 for another 24 hours and then treated with 2.5-20μM KU60019 (A8336, ApexBio) or 0.25-2μM AZD0156 (B7822, ApexBio) and a combination of 5-20μM fenofibrate (F6020-5G, Sigma) in RPMI-1640 supplemented with 0.1% FBS for 3 days. KU60019 and AZD0156 were administered 5 hours prior to fenofibrate. All drugs were supplied to the media daily. Colony formation was visualized by fixing cells in 1% paraformaldehyde for 5 min and staining with 0.05% crystal violet for 20 min. Wells were destained for 5 min in 500 mL 10% acetic acid. Absorbance (590nm) was measured using a spectrophotometer (Spectra Max 190). Each sample was assessed in triplicate. The synergy studies were further analyzed using Combenefit (36).

### Flow cytometry

Cells (5×10^6^/well in 12-well plates) were seeded, allowed to adhere overnight, and washed twice with 1x PBS the next day. Cells were cultured cell with serum-free RPMI-1640 for another 24 hours. Ovcar3 cells were then treated 1μM KU60019 and 10μM fenofibrate; Ovcar10 cells were treated 10μM KU60019 and 25μM fenofibrate in RPMI-1640 + 0.1% FBS for 3 days. KU60019 was administered 5 hours prior to fenofibrate. KU60019 and fenofibrate were supplied to the media daily. Cells were harvested by trypsin and washed twice with PBS. Cells were suspended in 0.5μg/ml 7-AAD (13-6993-T500, Tonbo Biosciences) in 1mL staining solution [900μL H2O + 100μL NaCitrate (380mM)] for 15 minutes at room temperature. Cells were run on a 10-color FACSCanto flow cytometer (BD Biosciences). Data were analyzed using FlowJo software (Ashland, OR).

### Western blotting

Cell lysates were collected in 1X sample buffer (2% SDS, 10% glycerol, 0.01% bromophenol blue, 62.5mM Tris, pH 6.8, 0.1M DTT) and boiled for 10 min at 95°C. Protein concentration was determined using the Bradford assay. Proteins were resolved using SDS-PAGE gels and transferred to nitrocellulose membranes (Fisher Scientific) (110mA for 2 h at 4°C). Membranes were blocked with 5% nonfat milk or 4% BSA in TBS containing 0.1% Tween-20 (TBS-T) for 1 h at room temperature. Membranes were incubated overnight at 4°C in primary antibodies (anti-lamin B1, ab16048, Abcam, 1:5000; anti-β-Actin, A1978, Sigma, 1:10,000) in 4% BSA/TBS + 0.025% sodium azide. Membranes were washed 4 times in TBS-T for 5 min at room temperature after which they were incubated with HRP-conjugated secondary antibodies (Cell Signaling, Danvers, MA) for 1 h at room temperature. After washing 4 times in TBS-T for 5 min at room temperature, proteins were visualized on film after incubation with SuperSignal West Pico PLUS Chemiluminescent Substrate (ThermoFisher, Waltham, MA).

### Immunofluorescence

Cells (5×10^6^/well in 12-well plates) were seeded overnight and washed with 1x PBS twice on next day. Cell were cultured with serum-free RPMI for another 24 hours. Ovcar3 cells were then treated 1μM KU60019 and 10μM fenofibrate; Ovcar10 cells were treated 10μM KU60019 and 25μM fenofibrate in RPMI + 0.1% FBS for 3 days. KU60019 was administered 5 hours prior to fenofibrate. KU60019 and fenofibrate were supplied daily. Cells were fixed in 4% paraformaldehyde for 10min and permeabilized in 0.2% Triton X-100 for 5 min. Cells then were stained with PML (1:200, Santa Cruz, Cat# sc-966) in 3% BSA/PBS at room temperature for 1 h. Cells were further incubated with 0.15 μg/ml DAPI in PBS (1 min), mounted, and sealed. Cells were washed three times and then incubated in Cy3 anti-mouse secondary antibody (1:5000, Jackson ImmunoResearch Labs, Cat# 715-165-150) in 3% BSA/PBS at room temperature for 1 h. Finally, cells were incubated with 0.15 μg/ml DAPI in PBS for 1min, washed three times with PBS, mounted, and sealed. Images were acquired at room temperature using a Nikon Eclipse 90i microscope with a 20x/0.17 objective (Nikon DIC N2 Plan Apo) equipped with a CoolSNAP Photometrics camera.

### Senescence-associated beta-galactosidase staining

SA-β-Gal staining was performed as previously described (37). Cells were fixed in 2% formaldehyde/0.2% glutaraldehyde in PBS for 5 min and stained (40 mM Na_2_HPO_4_, 150 mM NaCl, 2 mM MgCl_2_, 5 mM K_3_Fe(CN)_6_, 5 mM K_4_Fe(CN)_6_, and 1 mg/ml X-gal) overnight at 37 °C in a non-CO_2_ incubator. Images were acquired at room temperature using an inverted microscope (Nikon Eclipse Ts2) with a 20X/0.40 objective (Nikon LWD) equipped with a camera (Nikon DS-Fi3). Each sample was assessed in triplicate and at least 100 cells per well were counted (> 300 cells per experiment).

### Quantification and Statistical Analysis

GraphPad Prism version 8.0 was used to perform statistical analysis. One-way ANOVA or t-test were used as appropriate to determine p-values of raw data. p-values < 0.05 were considered significant.

## Results

### ATM is wildtype and upregulated in HGSOC

ATM is thought to be a tumor suppressor as mutations in ATM predispose patients to tumorigenesis (7,8). However, recent data suggest that ATM expression is required for malignant transformation in response to oncogenic stress (11), suggesting that ATM plays a context-dependent role in tumorigenesis. Analysis of TCGA data demonstrate that the proportion of patients with ATM mutations varies widely among different tumor types (**Fig. 1A**). Interestingly, although HGSOC is known to have defects in HR (4,38), ATM is only mutated in ~0.5-2% of these patients (**Fig. 1B**). Compared to normal fallopian tube, the likely cell-of-origin for HGSOC (39,40), HGSOC specimens do not have markedly increased ATM protein expression (**Fig. 1C**) (41). However, phosphoproteomics analysis demonstrates that ATM kinase activity is significantly upregulated in HGSOC compared to normal fallopian tube, as indicated by increased S1981 autophosphorylation (**Fig. 1D**). Previous reports have shown that high ATM expression at both the protein and mRNA level is associated with worse overall and progression-free survival (12). Together, these data suggest that ATM is wildtype and its signaling pathway is upregulated in HGSOC, indicating that it may be an actionable target for these patients.

**Figure 1.**
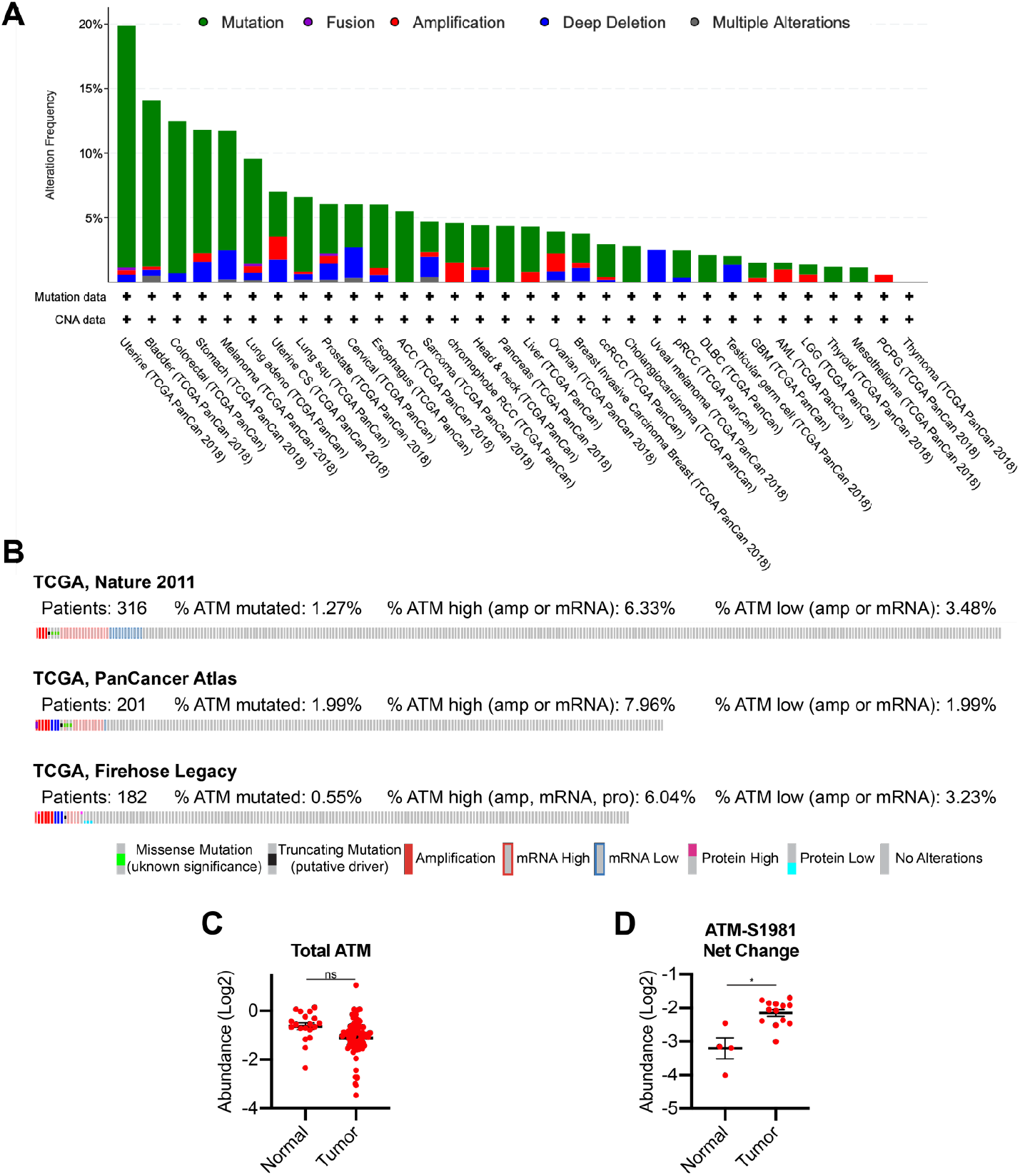
ATM is wildtype and upregulated in high grade serous ovarian cancer patients. **(A)** Analysis of ATM alterations in TCGA PanCancer Atlas Studies. **(B)** Analysis of ATM alterations in TCGA HGSOC studies. **(C)** Total ATM protein expression in normal fallopian tube and HGSOC specimens. ns = not significant. **(D)** Phosphorylated ATM (S1981) protein expression in normal fallopian tube and HGSOC specimens. *p<0.001

### ATM low HGSOC tumors display a metabolic gene signature that is targetable

ATM may be an actionable target in HGSOC. However, many preclinical studies have demonstrated that inhibition of ATM as a single agent is not likely to be effective (14–18).

However, combined inhibition of ATM and other therapeutic agents has shown potential to inhibit cancer cell survival both *in vitro* and *in vivo* (13,14,18,42). We reasoned that genes and pathways that are inversely correlated with ATM expression may lead to the identification of novel targets for combinatorial therapeutics. Gene set enrichment analysis (GSEA) using HGSOC patient data from TCGA suggested that terms related to both cell cycle and DNA damage are enriched in ATM-low tumors as expected (**Table S1**). Interestingly, we also found that metabolic pathways such as oxidative phosphorylation, MYC signaling, mTORC1 signaling, fatty acid metabolism, glycolysis, and TCA cycle metabolism are enriched in ATM-low HGSOCs, suggesting that target these pathways may be synergistic with ATM inhibitors (**Fig. 2A and Table S1**). To further investigate potential drug combinations, we took advantage of the Dependency Map database (depmap.org) PRISM drug repurposing screen (32). We divided the cell lines into ATM high and ATM low based on protein expression, and looked for FDA-approved drugs that kill ATM-low cells to a greater degree (log_2_FC<0; FDR<25%). We found 17 “hits” that met these criteria (**Fig. 2B and Table 1**). Of these, the majority were inhibitors of EGFR, which have already been shown to have synergistic effects with ATM inhibitors (43). Interestingly, fenofibrate was found as one of the hits (**Fig. 2B-C and Table 1**). Fenofibrate is a PPARα agonist that has previously been shown to inhibit multiple tumor-promoting metabolic pathways including oxidative phosphorylation and glycolysis (27,44). These data indicate that inhibition of metabolic pathways using the FDA-approved drug fenofibrate may be a novel therapy to use in combination with ATM inhibitors.

**Figure 2.**
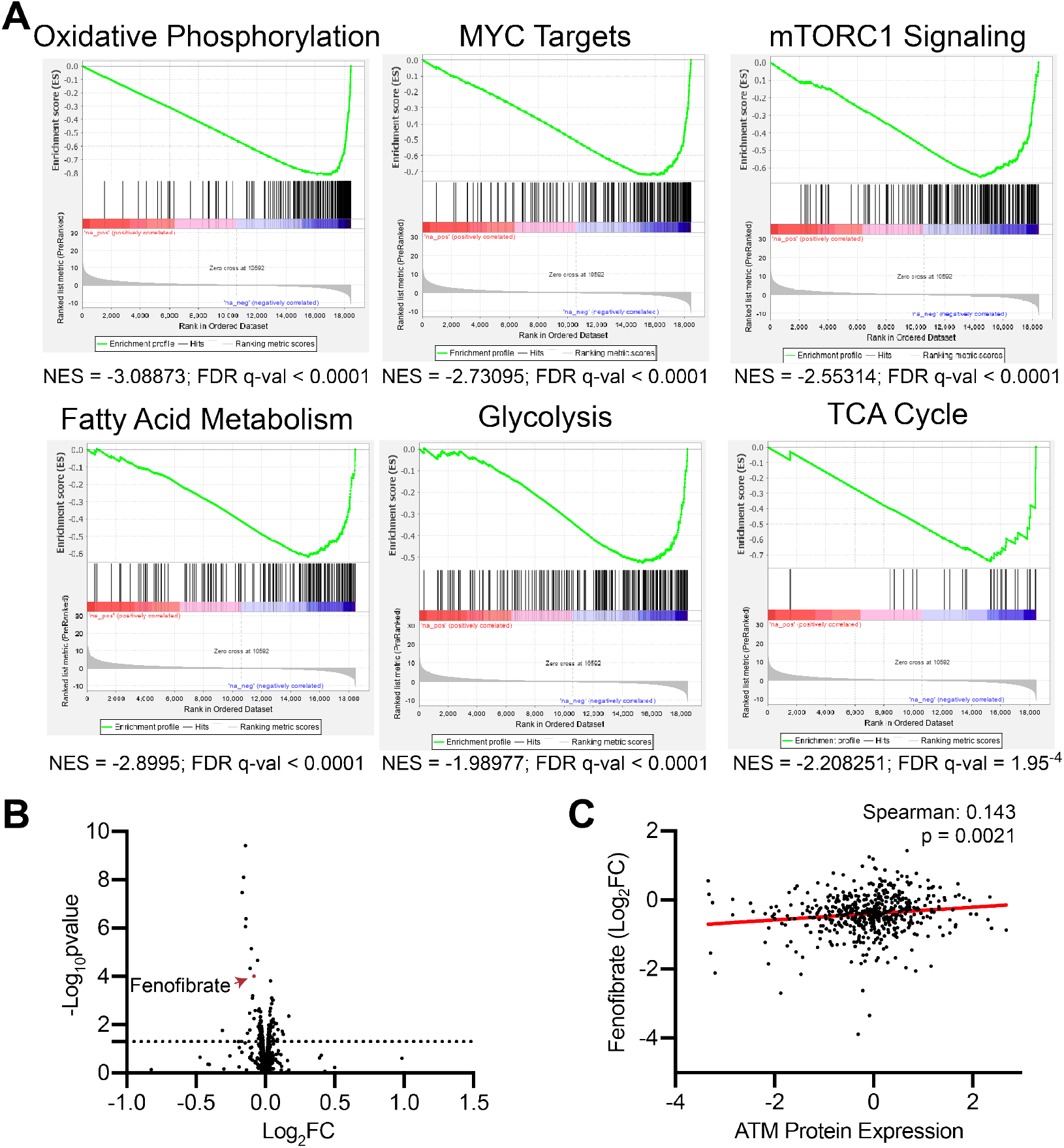
ATM-low specimens have increased metabolic gene signature and are more sensitive to fenofibrate. **(A)** GSEA enrichment analysis in ATM-low vs. ATM-high HGSOC specimens from TCGA. **(B)** Volcano plot of FDA-approved drugs in ATM-low vs. ATM-high cell lines. Line represents p-value<0.05. **(C)** Correlation between fenofibrate sensitivity and ATM protein expression. Note that lower Log_2_FC = greater sensitivity.

**Table 1:** FDA-approved drugs that sensitize ATM-low cells to a greater extent than ATM-high cells.

| Drug Name | Target | Log<sub>2</sub>FC<br>(ATM-low<br>vs. ATM-high) | FDR |
| --- | --- | --- | --- |
| lapatinib | EGFR,<br>ERBB2 | -0.1431668 | 4.54E-07 |
| osimertinib | EGFR | -0.1575673 | 6.22E-06 |
| ibrutinib | BTK | -0.1669926 | 1.95E-05 |
| gefitinib | EGFR | -0.139171 | 0.00012 |
| afatinib | EGFR,<br>ERBB2,<br>ERBB4 | -0.1432169 | 0.000225 |
| icotinib | EGFR | -0.1019151 | 0.001532 |
| olmutinib | EGFR | -0.0559589 | 0.003793 |
| vandetanib | EGFR,<br>VEGFR,<br>RET | -0.1091835 | 0.007295 |
| fenofibrate | PPARA | -0.0821997 | 0.012972 |
| spironolactone | Various | -0.0930345 | 0.067266 |
| erlotinib | EGFR | -0.0937315 | 0.075598 |
| alfacalcidol | VDR | -0.0450776 | 0.177706 |
| bosutinib | Bcr-Abl,<br>Src | -0.0724218 | 0.189293 |
| maxacalcitol | VDR | -0.0500369 | 0.189293 |
| paricalcitol | VDR | -0.0490078 | 0.191315 |
| loperamide | Various | -0.0341769 | 0.19634 |
| dequalinium | Various | -0.1146281 | 0.219346 |

### PPARα is associated with ATM expression

ATM-low cancer cells are slightly, although significantly more sensitive to fenofibrate (**Fig. 2**), suggesting that these cells may have low PPARα signaling. Indeed, there is a positive correlation between ATM and PPARα in both cell lines and HGSOC patient samples (**Fig. 3A-B**), suggesting that ATM-low cancers have low PPARα, which may be therapeutically exploited using fenofibrate.

**Figure 3.**
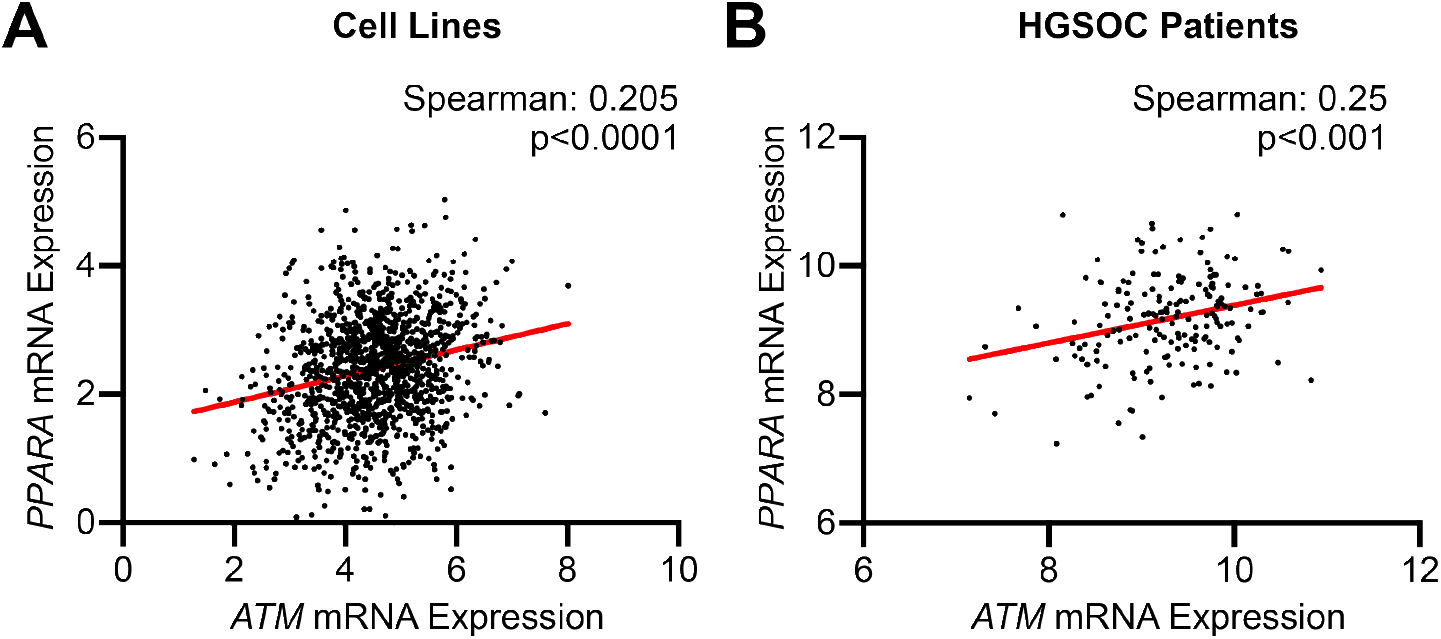
PPARα expression is associated with ATM expression in cell lines and HGSOC patient specimens. **(A)** Correlation between *ATM* and *PPARA* (encoding PPARα) expression in cell lines from Dependency Map. **(B)** Correlation between of *ATM* and *PPARA* expression in HGSOC patient specimens from TCGA.

### Combination of ATM inhibitor and fenofibrate is synergistic in HGSOC cells

We found that ATM-low cancer cells are more sensitive to fenofibrate (**Fig. 2**) and HGSOC cells with low ATM expression have low PPARα signaling (**Fig. 3**), suggesting that the combination of fenofibrate with ATM inhibitors may be synergistic in HGSOC cells. Indeed, inhibition of ATM using two small molecule inhibitors (KU60019 and AZD0156) synergized with fenofibrate in multiple HGSOC cell lines to decrease cell proliferation (**Fig. 4A-B**). While cell death, as determined by 7AAD staining, was not affected (**Fig. 4C**), we observed multiple senescence markers, including increased SA-β-Gal and PML bodies and decreased lamin B1 expression in the combination treated cells (**Fig. 4D-E**). Together, these data demonstrate that the combination of ATM inhibition and fenofibrate treatment is synergistic in HGSOC through induction of senescence.

**Figure 4.**
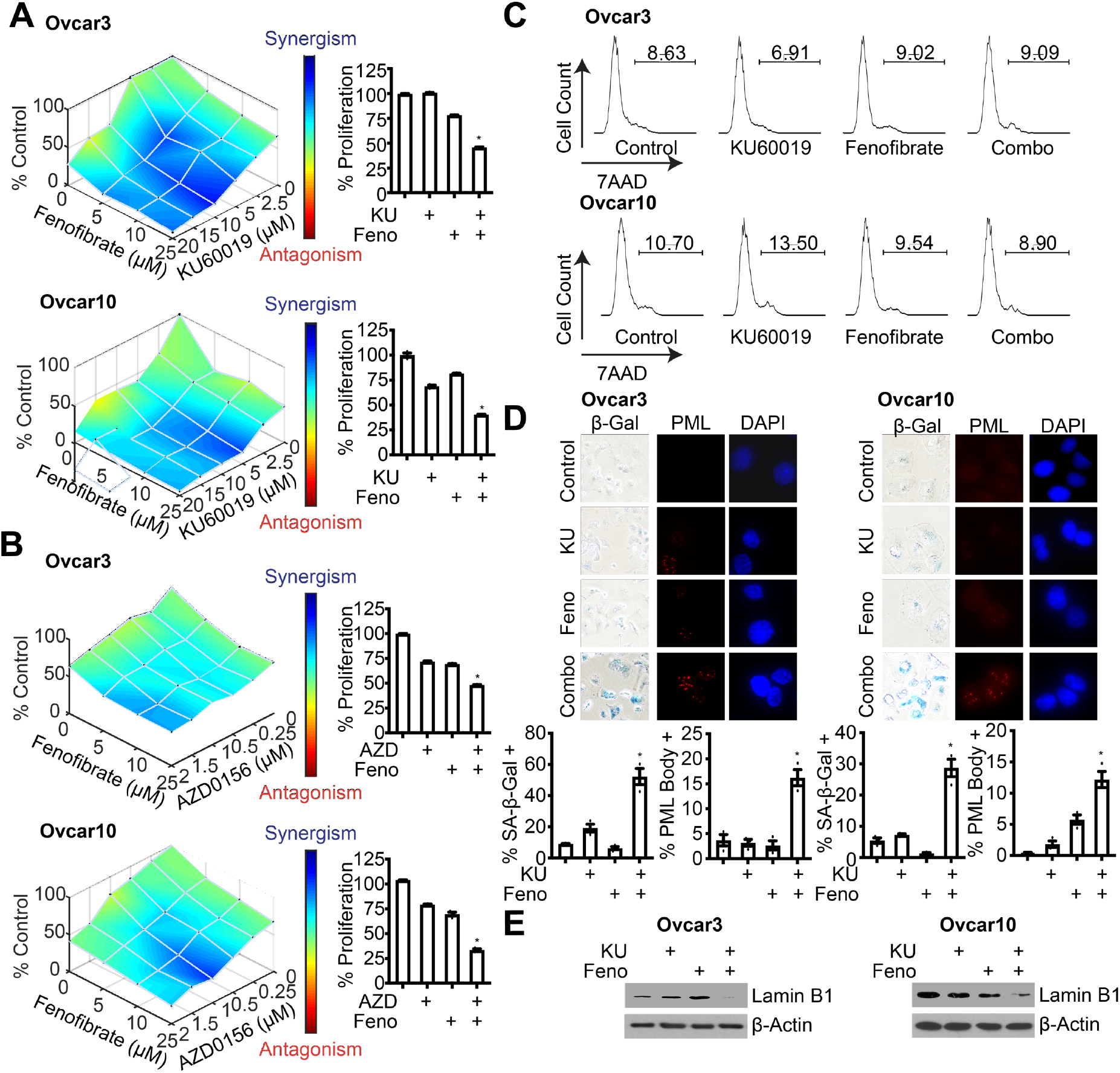
Combined inhibition of ATM and treatment with fenofibrate is synergistic in HGSOC cell lines. **(A-B)** Ovcar3 and Ovcar10 cells were treated with the ATM inhibitors KU60019 (A) or AZD0156 (B) or the PPARα agonist fenofibrate alone or in combination for 3 days. Proliferation was determined by crystal violet staining. n=3/group, one of 3 experiments is shown. *p<0.001 **(C)** Cell death was assessed by 7AAD staining. n=3/group, one of 3 experiments is shown. Data represent mean ± SEM. **(D)** SA-β-Gal and PML body staining. n=3/group, one of 3 experiments is shown. Data represent mean ± SEM .*p<0.001. **(E)** Immunoblot analysis of lamin B1. β-actin was used as a loading control. One of 3 experiments is shown.

## Discussion

Many tumors are addicted to DNA repair signaling (45), which has led to investigation of ATM inhibitors as cancer therapies (13,14). These inhibitors are not effective as a monotherapy, demonstrating the need to identify new targets for combinatorial therapeutic strategies. Here, we identified fenofibrate as a potential drug combination for use with ATM inhibitors. This may be due in part to upregulation of multiple metabolic pathways in ATM-low cancers. The combination induced senescence, a stable cell cycle arrest that is considered a positive patient outcome (46–48). Together, our results provide rationale for exploration of drugs that modify metabolism as combinatorial therapies with ATM inhibitors.

ATM is an critical mediator of DNA DSB repair through HR (7,8,14). Based on this, and the fact that both A-T patients and Atm knockout mice are predisposed to cancer, it has been well-appreciated that ATM is a tumor suppressor (7,8,10). However, many cancer cells are addicted to DNA damage repair and ATM signaling, and a recent study demonstrated that ATM is required for breast tumorigenesis (11). This suggests that in some contexts, ATM may act as an oncogene. Indeed, we found that HGSOC patients overwhelmingly harbor wildtype ATM alleles, and ATM kinase activity is upregulated in HGSOC samples compared to fallopian tube (**Fig. 1**). This is consistent with another paper that demonstrated increased ATM nuclear expression in serous ovarian cancer, which was associated with worse survival (12). Therefore, in the context of the ~50% of HGSOCs with HR-proficient disease, ATM can be considered to act more like an oncogene than tumor suppressor. This suggests that inhibition of ATM may be relevant for a large subset of HGSOC patients. As these patients often have worse survival that HR-deficient patients, identification of novel therapies is a clinical need. Indeed, ATM inhibitors, while not effective on their own, have shown promise in multiple cancer cell types in combination with irradiation or other DNA damage agents (13,14,16,18).

ATM has multiple functions outside of its role in DNA repair (8,19–21,49). We found that ATM-low HGSOC specimens showed multiple terms related to metabolism, including oxidative phosphorylation, TCA cycle metabolism, fatty acid metabolism, glycolysis, and signaling related to both MYC and mTORC1 (**Fig. 2**). This is consistent with previous reports from us and others that have shown suppression of ATM alters metabolic functions in multiple ways (8,19–21,50–54). For instance, we found that inhibition of ATM increases consumption of multiple metabolites, including glucose, glutamine, and branched chain amino acids (19,20,49). Similarly, A-T patients and A-T patient cells display multiple metabolic phenotypes, including mitochondrial dysfunction, insulin resistance, and an increased susceptibility to both diabetes and cardiovascular disease (7,8). Considering ATM inhibitors are not effective as a monotherapy, exploring metabolic vulnerabilities of ATM inhibited cancer cells may lead to additional combinatorial therapeutic strategies.

PPARα has not been well-studied in ovarian cancer. One study found that PPARα is expressed at a higher level in pleural effusions than in primary or metastatic tumors (55). Interestingly, they also found PPARα to be expressed at a much lower level than either PPARδ or PPARγ, suggesting that PPARα signaling while present may be low in ovarian cancers. Consistently, multiple studies have found that PPARα agonists inhibit ovarian cancer proliferation and growth both *in vitro* and *in vivo* through a variety of mechanisms (56–58). We also found that at the dose and timing used in this study, the PPARα agonist fenofibrate moderately inhibits HGSOC cell proliferation (**Fig. 4**). Fenofibrate is an FDA-approved with limited toxicity, suggesting that further studies are warranted to determine whether PPARα agonists hold promise for HGSOC therapy.

Here, we found that the combination of ATM inhibition and fenofibrate is synergistic by inducing senescence (**Fig. 4**). Senescence is a tumor suppressive mechanism due to its inhibition of cancer cell proliferation (47,48,59). Recent data from HGSOC patients suggest that senescence occurs *in vivo* after therapy and is associated with a better outcome (46). Indeed, other publications have also indicated that senescence is a beneficial therapeutic response (47,48,60–63). The mechanism of senescence induction by the combination of ATM inhibition and fenofibrate remains to be explored. We found that PPARα expression is associated with ATM expression both in HGSOC patient samples (**Fig. 3**), and ATM-low patients have altered metabolic pathways (**Fig. 2**). Thus, is possible that enhanced PPARα signaling using an agonist competes with metabolic pathways that are altered in ATM-low/inhibited cells. For instance, we previously published that ATM suppression increased glucose uptake and utilization (19), whereas fenofibrate decreases this process (27). Similarly, others have reported that Atm knockout cells display mitochondrial dysfunction (54). This may be further exacerbated by fenofibrate, which decreases mitochondrial metabolism and oxidative phosphorylation through a variety of mechanisms (27,44). Regardless of the mechanism, given that fenofibrate is FDA-approved and has an excellent safety profile, our results provide a proof-of-principle study to combine fenofibrate or other PPARα agonists with ATM inhibitors. Our studies may also suggest that cancers with a high prevalence of ATM mutations (for instance melanoma) may be especially sensitive to PPARα agonists.

## Conclusions

The present study shows that ATM is wildtype and upregulated in HGSOC, which corresponds to low PPARα expression. ATM-low cells display changes in multiple metabolic pathways that reveal a therapeutic vulnerability to use for combinatorial treatment with ATM inhibitors. The combined inhibition of ATM and treatment with the PPARα agonist fenofibrate was synergistic in HGSOC cell lines. Thus, our study provides a new potential combination therapy for HGSOCs that are HR-proficient. As multiple metabolic terms were associated with ATM-low HGSOC specimens, we predict that additional metabolic drugs may have synergistic effects with ATM inhibitors.

## List of Abbreviations

EOC: Epithelial ovarian cancer
HGSOC: high grade serous ovarian cancer
HR: homologous recombination
TCGA: The Cancer Genome Atlas
GSEA: Gene Set Enrichment Analysis
ATM: Ataxia Telangiectasia Mutated
PARP: poly(ADP)ribose polymerase
DSB: DNA double strand break
PPAR: Peroxisome Proliferator Activated Receptor

## Declarations

### Availability of data and materials

All data generated or analyzed during this study are included in this published article

### Authors’ contributions

CWC and KMA are responsible for conception of the study. CWC, ESD, RB, and KEL performed experiments, collected data, and participated in data analysis and interpretation. CWC and KMA drafted the manuscript. All authors read and approved the final manuscript.

